# Transposable elements contribute to dynamic genome content in maize

**DOI:** 10.1101/547398

**Authors:** Sarah N Anderson, Michelle C Stitzer, Alex B. Brohammer, Peng Zhou, Jaclyn M Noshay, Cory D. Hirsch, Jeffrey Ross-Ibarra, Candice N. Hirsch, Nathan M Springer

## Abstract

Transposable elements (TEs) are ubiquitous components of eukaryotic genomes and can create variation in genomic organization. The majority of maize genomes are composed of TEs. We developed an approach to define shared and variable TE insertions across genome assemblies and applied this method to four maize genomes (B73, W22, Mo17, and PH207). Among these genomes we identified 1.6 Gb of variable TE sequence representing a combination of recent TE movement and deletion of previously existing TEs. Although recent TE movement only accounted for a portion of the TE variability, we identified 4,737 TEs unique to one genome with defined insertion sites in all other genomes. Variable TEs are found for all superfamilies and are distributed across the genome, including in regions of recent shared ancestry among individuals. There are 2,380 genes annotated in the B73 genome located within variable TEs, providing evidence for the role of TEs in contributing to the substantial differences in gene content among these genotypes. The large scope of TE variation present in this limited sample of temperate maize genomes highlights the major contribution of TEs in driving variation in genome organization and gene content.

**Significance Statement:** The majority of the maize genome is comprised of transposable elements (TEs) that have the potential to create genomic variation within species. We developed a method to identify shared and non-shared TEs using whole genome assemblies of four maize inbred lines. Variable TEs are found throughout the maize genome and in comparisons of any two genomes we find ~20% of the genome is due to non-shared TEs. Several thousand maize genes are found within TEs that are variable across lines, highlighting the contribution of TEs to gene content variation. This study creates a comprehensive resource for genomic studies of TE variability among four maize genomes, which will enable studies on the consequences of variable TEs on genome function.

## Introduction

In many eukaryotic genomes, genes comprise less than 5% of the genome. The remaining sequence includes low-copy intergenic sequences as well as repetitive sequences including tandem repeats and transposable elements (TEs). TEs were first identified in maize through Barbara McClintock’s studies (1). Since then, they have been widely used as tools for genetic analysis (2–4) and have been shown to contribute to phenotypic variation. Depending on the method used to annotate TEs, 65% (structurally intact TEs) or 85% (all TE fragments) of the ~2.3Gb reference B73 maize genome is annotated as TEs (5, 6). TEs include several distinct classes of elements that have different structural features and mechanisms for replication (7). Retrotransposons (class I) replicate through an RNA intermediate and can be separated into long terminal repeat (LTR) retrotransposons and non-LTR retrotransposons (long interspersed elements (LINEs) and short interspersed elements (SINEs)). DNA transposons (class II) replicate through a DNA intermediate and can be divided into two orders: terminal inverted repeat (TIR) transposons and Helitrons. Within each of these orders there are families of elements that are classified based on sequence similarity, and the members of a family are likely mobilized by the same factors. These families can vary in size from a single member to >10,000 members (6).

TEs can have a variety of influences on gene function (8–11). These can include insertional mutagenesis as well as complex influences on gene expression through insertion into regulatory sequences or by influencing local chromatin (8, 11, 12). The transposon content of different individuals of the same species can vary substantially (13–15). Researchers have used PCR based techniques, such as TE display, to amplify fragments from the ends of TEs to demonstrate that TEs can be used as polymorphic markers in many species (16–19). Targeted sequencing of BAC clones for multiple haplotypes at several maize loci revealed that while gene content is largely conserved there are very different locus-specific TE insertions (20–22). At the *Bz* locus only 25-84% of the region includes alignable sequence when multiple haplotypes were compared. The non-alignable sequences were largely distinct TE insertions among the haplotypes (20, 21). Studies of TE variability in plants on a genome-wide scale have largely focused on using short-read resequencing data to map new insertions (23–27), however this approach can be complicated by non-alignable sequences and the repetitive nature of TEs.

The analysis of genic and other low-copy regions of the maize pan-genome has revealed extensive structural variation (28–33). The availability of multiple *de novo* genome assemblies for maize (6, 29, 34, 35) provides new opportunities to characterize the shared and polymorphic nature of TEs in the maize genome. Here, we implement a novel method to compare shared and variable TEs across homologous and co-linear blocks of maize genome assemblies to identify extensive TE variation among inbred lines. This variation represents all orders of TEs and is found throughout the genome, including in regions of recent shared ancestry.

Remarkably, we find that over half of B73 genes are near TEs that are variable across these lines. We present evidence for a substantial contribution of variable TEs to gene content differences among lines, with over 2,000 genes annotated in TEs present in B73, but absent in another line. Together, this highlights the important role of TEs in creating genome content variation in maize as well as potential impacts on transcriptional and phenotypic variation.

## Results

### Comparable structural annotations reveal similar TE content among genomes

Whole-genome assemblies of maize genotypes B73, W22, PH207, and Mo17 are available (6, 29, 34, 35). In order to compare the TE content in these genomes it is important to have a set of consistent structural annotations of TEs and TE families in each genome. Consistent annotations were performed with nesTEd (see methods for details), which is a modified version of the structural annotation approach previously used for annotation of TEs in the B73v4 and W22 reference genome assemblies (6, 34) that has been updated for improved annotation of TIR elements. Transposable elements are classified into families based on sequence similarity, superfamilies based on structural similarities, orders based on replication mechanism, and classes based on presence or absence of an RNA intermediate (7). In these annotations, nomenclature is consistent across the genome assemblies down to the TE family level, however individual elements do not have consistent unique IDs across genomes. The majority of DNA sequence (56% – 62%) for all genome assemblies are composed of LTR retrotransposons, with Helitrons and TIRs contributing approximately 4% and 3% of genomic sequence, respectively (Figure S1A, Table 1). The size of TE families varies, with some families containing only a single element per genome while others have > 1,000 insertions (Table S1). These annotations served as the basis for studying TE insertion variability across the four genomes.

**Table 1:** Annotated elements of each TE superfamily in maize genomes

|  | **TE Orders (Code)** | **Superfamily (Code)** | **B73** | **W22** | **Mo17** | **PH207** |
| --- | --- | --- | --- | --- | --- | --- |
| Class I<br>Retrotransposons | LTR retrotransposons (RL) | Copia (RLC) | 46,257 | 45,701 | 44,315 | 26,753 |
|  |  | Gypsy (RLG) | 75,503 | 71,774 | 70,144 | 48,561 |
|  |  | Unclassified LTR (RLX) | 20,430 | 18,958 | 23,303 | 18,210 |
|  | LINES (RI) | L1 (RIL) | 477 | 459 | 439 | 436 |
|  |  | RTE (RIT) | 296 | 284 | 295 | 243 |
|  | SINES (RS) | (RST) | 892 | 907 | 830 | 654 |
| Class II<br>DNA transposons | TIR transposons (DT) | hAT (DTA) | 5,087 | 5,099 | 5,153 | 2,225 |
|  |  | CACTA (DTC) | 2,768 | 2,553 | 2,776 | 718 |
|  |  | Pif/Harbinger (DTH) | 63,201 | 57,879 | 46,020 | 49,837 |
|  |  | Mutator (DTM) | 927 | 885 | 950 | 677 |
|  |  | Tc1/Mariner (DTT) | 66,168 | 60,073 | 57,559 | 55,623 |
|  |  | Unclassified TIR (DTX) | 34,689 | 28,425 | 27,681 | 22,563 |
|  | Helitrons (DH) | Helitron (DHH) | 21,529 | 21,416 | 22,257 | 17,517 |

It is important to note that a comparison of the TE content among the genomes solely based upon the annotations of intact elements will contain a number of false-positive calls of polymorphisms. The nesTEd annotation defines TEs based on the structural features specific to each superfamily. Given the reliance on short sequence signatures and the choice to only include structurally intact elements, annotated elements represent a subset of the true TE-derived sequences of the genome as evident by the genomic space that is RepeatMasked but not annotated as TEs (Figure S1). The stringent requirement for intact structural features can result in missed annotations resulting from biological noise such as single nucleotide changes in the target site duplications (TSD) or from technical noise such as N’s in the assembly. This results in many examples where a TE is present in two genotypes based on the homology of the region, but the TE is only annotated in one of the genomes (Figure S1). Therefore, we developed an approach to utilize the TE annotations as a starting point to document shared and non-shared TE insertions relative to other genomes, but that does not require the element to be annotated in both genomes.

### Substantial TE variation is found across genomes

Classification of specific TE insertions as shared (present in both genotypes at a specific location) or non-shared (polymorphic) between genotypes is complicated by the highly repetitive nature of TEs. In order to reduce the complexity of the problem we developed a robust two step approach to search for shared/non-shared TE insertions within windows anchored by collinear genes (Figure 1). Alignments of windows were parsed to classify shared (TE.1), non-shared (TE.2), and unresolved (TE.3) elements based on homology of sequences annotated as transposable elements. For a subset of elements, homology for the flanking sequences (200bp) can be used to define coordinates for shared elements (hereafter referred to as shared site defined, TE.4) and TE empty sites (hereafter referred to as non-shared site defined, TE.5). This results in the classification of shared and non-shared elements throughout the genome with a subset of these with highly similar flanking sequence that can be site-defined. The non-shared site defined elements represent recent insertions in shared sequences or precise excision events while the remaining non-shared elements for which the flanking sequences are not similar reflect large deletions or insertions into non-shared sequences. This approach was implemented to compare annotated TEs in each of the four genomes to the other three genomes in a pairwise fashion in order to classify shared or non-shared TE insertions. Importantly, we noted that across all contrasts, 5.1% of shared elements did not overlap any TE annotation in the target genome, highlighting the importance of using more nuanced homology based approaches beyond simple comparison of annotations.

**Figure 1:**
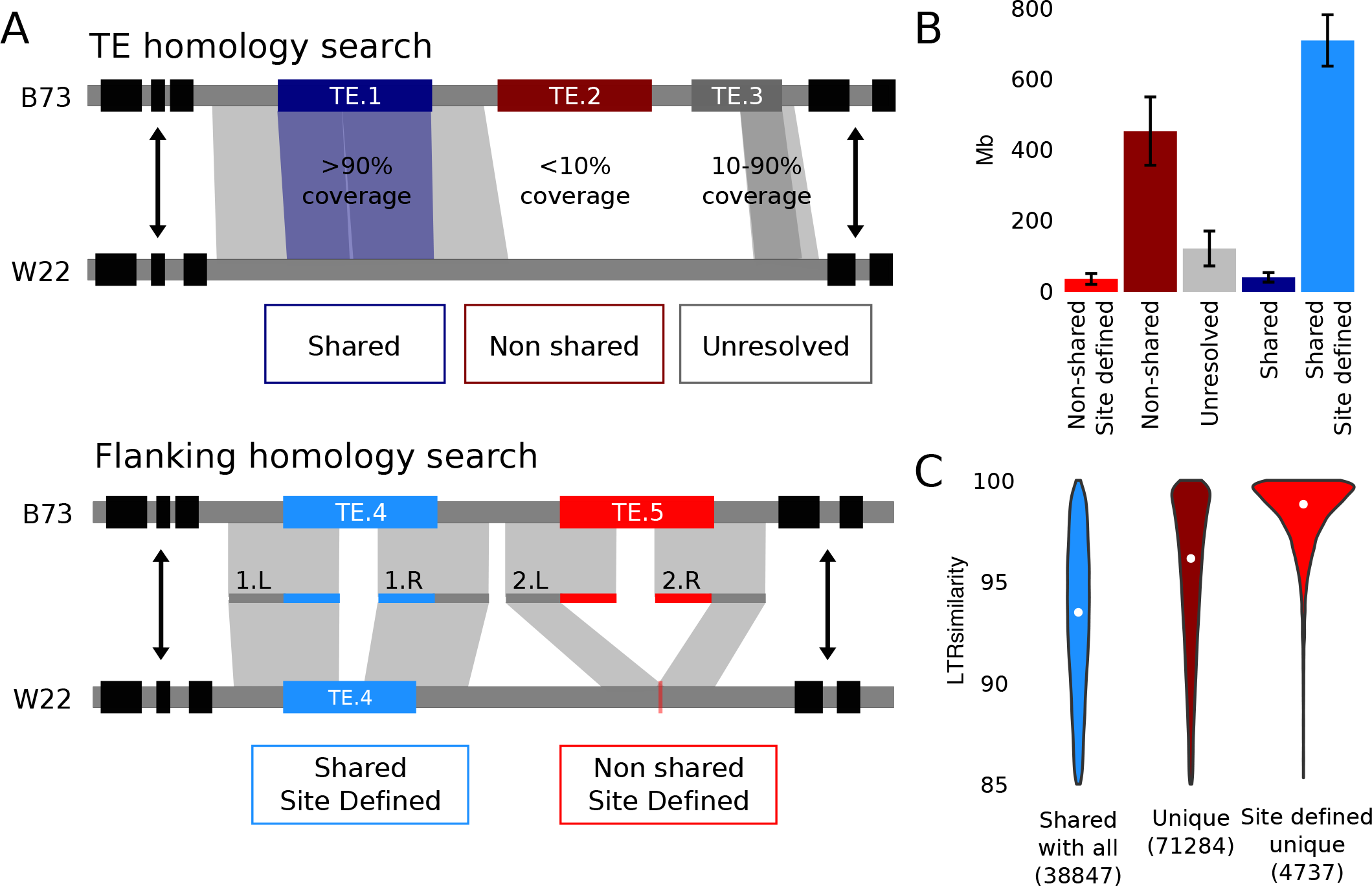
Comparison of shared and variable TE insertions from whole genome assemblies. A. Schematic representation of our method to define shared and variable TEs between assembled genomes. The TE homology search parses alignments to define shared (dark blue), non-shared (dark red), and unresolved (gray) TEs and the flaking homology search utilizes alignments of 200 bp flanking sequences to define coordinates for site-defined shared (blue) and non-shared (red) TEs. B. The average genomic space of TEs with each assignment across all 12 pairwise contrasts is shown (error bars represent +/− the standard deviation). C. The distribution of LTR similarity for LTRs shared across all four genomes (blue) as well as TEs unique to one genome. The non-shared elements that are unique to only one genome were separated into TEs that are site defined in every contrast (red) and TEs that are classified as missing based on lack of homology for the TE region in at least one contrast (dark red). LTR similarity is used as an approximation for TE age, with high similarity for young insertions and low similarity denoting old insertions. The number of LTR elements in each category are listed in parenthesis, and the median value is marked with a white point.

In total, pairwise contrasts between genomes reveals an average of ~500 Mb of non-shared TE sequences, representing ~20% of the genome content that is variable between any combination of these inbred lines (Figure 1B). The vast majority of shared elements are site-defined, indicating high sequence similarity at the flanking region in addition to the TE. In contrast, only ~5% of non-shared calls are site defined. Across all four genomes, we found 46,122 LTR, 19,744 TIR, and 2,529 Helitron non-shared site defined calls (Table S2). We evaluated several loci that correspond to sequenced and manually annotated BACs (20, 21). We found our genome-wide approach was consistent with prior calls for those TEs that are annotated in both approaches (Figure S2). We predicted that non-shared site defined TEs often represent new insertions in otherwise shared haplotypes, while the non-shared TEs defined by lack of homology would include a larger range of TE ages since this pattern could result from a combination of new insertions, deletions, and haplotype diversity. To support this, we assessed TE age of LTR retrotransposons, which have identical LTR sequences upon insertion that diverge over time. We found that non-shared site defined TEs unique to one genome were enriched for younger insertions than shared TEs or those unique but not site defined (Figure 1C).

### Monitoring levels of TE variability across superfamilies and families

A non-redundant set of TEs was developed from the subset of TEs that could be resolved as shared or non-shared across all contrasts. Across all four assembled genomes 457 Mb of TEs (110,643 elements) are shared in all genomes. Another 1.6 Gb (399,890 elements or 78% of assessed TEs) are variable TEs missing in at least one of the genomes. On average each genome has 207 Mb (49,571 elements) that are unique to that genome (Figure 2A). Different superfamilies of TEs show subtle differences in distributions across these four genomes, particularly among TIRs which range from 75.5% variable (DTT, Mariner) to 86.4% variable (DTM, Mutator) (Figure S3A). In all cases, there are fewer elements that are unique to PH207 relative to other inbreds, which likely reflects variable assembly quality rather than biological differences.

**Figure 2:**
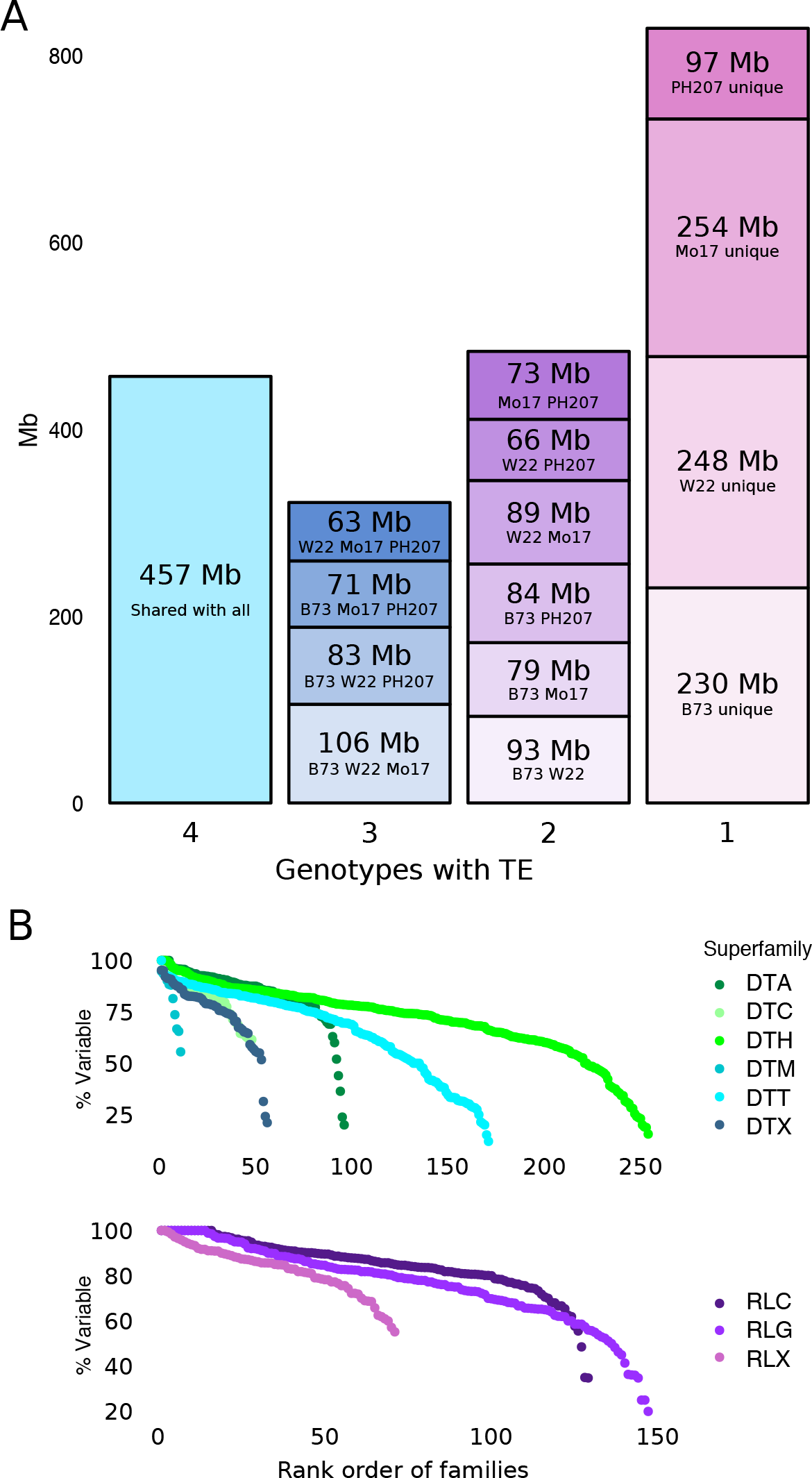
Shared and non-shared TE content among four maize inbred lines. A. All non-redundant TEs that could be fully resolved as shared or non-shared in all contrasts were classified based on the number of genomes containing the elements. The amount genome sequence shared in each combination of genotypes is listed. B. For families with at least 20 members in the non-redundant TE set the proportion of members that exhibit polymorphic presence in the four genomes was determined. Within each superfamily the proportion of elements that are polymorphic were used to generate a rank order from the most to least variable.

While the level of TE variability among the four genomes is similar across orders and superfamilies of TEs, there are many specific TE families that show highly biased distributions. The analysis of TIR and LTR families with at least 20 members across the four inbreds reveals that some families are highly conserved, while others have >75% variable members (Figure 2B, S3B). The highly variable TE families include examples in which there are members of the family in each genome at different loci as well as cases in which one genome has many more family members than the others (Figure S4A). In many cases, the non-shared copies are distributed throughout the genome rather than clustered (Figure S4B). TE families that are unique to one of the four genomes represent the most extreme type of variation. There are 7,576 TE families that are unique to one of the four genomes. The vast majority (96%) of these families contain a single element. However, there were 15 examples of inbred-specific TE families with at least 4 members in the resolved TE set (Table S3). These unique families likely represent a combination of new TE movement in one lineage and dying TE families that have been lost from most haplotypes.

### Variable TE insertions in maize are found genome-wide

The genomic distribution of TE variability was investigated by looking at the distribution of B73 TEs that are shared and variable (Figure 3A, S5A). Variable TEs are found across chromosomes, with regions of high and low variability located on chromosome arms and in paracentromeric regions (Figure 3, S5). The analysis of gene density within 1 Mb windows of the genome reveals a relationship between local gene density and TE variation levels (Figure 3B). The regions of the genome with the lowest gene density exhibit a wide range of TE variation levels while regions with higher gene density tend to have higher levels of TE variation (Figure 3B). Similarly, local levels of CHG methylation are related to TE variability such that the most highly methylated regions of the genome include regions with the least TE variability (Figure S5C). Prior studies have found evidence for biased gene fractionation in the two sub-genomes of maize that resulted from an allopolyploid fusion event (36–38). However, there was no difference in the level of variable TEs within the two sub-genomes of maize (Figure S5D).

**Figure 3:**
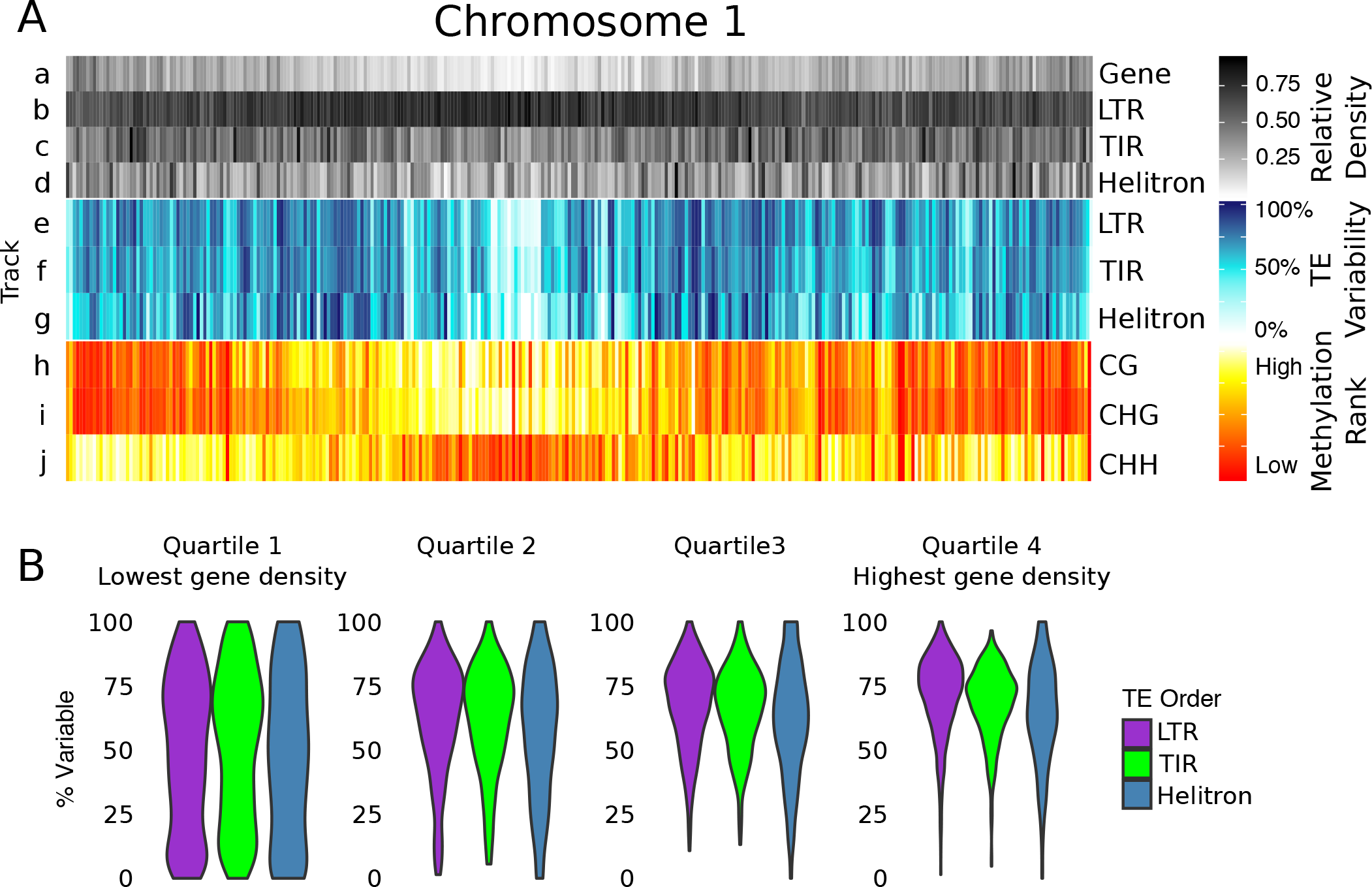
Genomic distribution of shared and variable TEs. A. The distribution of gene density, TE variability and DNA methylation in 1 Mb bins along B73 chromosome 1. Tracks a-d show the relative gene density and TE density for LTRs, TIRs, and Helitron. Tracks e-g show the proportion of B73 TEs that are variable in each bin for LTRs, TIRs and Helitrons, and tracks h-j show the rank order of methylation in the CG, CHG, and CHH context. B. The gene density in each 1 Mb bin was used to splits the bins into four quartiles from lowest to highest gene density. The proportion of variable TEs within each bin were determined for each superfamily.

The four genotypes used in this study are all temperate maize germplasm that contain regions of identity by state (IBS). These regions provide a unique opportunity to look for new transposition events as indicated by their variation within regions of otherwise complete sequence similarity (Figure 4). In total there are 235.25 Mb of IBS regions between B73 and the other three genomes across 37.64 cM (Tables S4, S5) and, as expected, these regions largely include shared TEs (Table S4). However, there are a set of non-shared TEs in these regions that reflect new insertion events, deletions that remove the TEs, or gaps in the assemblies. There are 29 examples of site-defined polymorphic TEs in 19 families, including 8 TIR elements and 21 LTR elements that likely represent new TE insertion events in these IBS regions. Since TEs require expression for movement, we assessed the level of expression for these TE families across 23 B73 tissues (39). Expression was detected for 14 of these families (74%, while only 13% of all families are expressed), with some showing expression across all tissues and others showing higher expression in reproductive tissues (Figure S6A). The ability to estimate ages for LTR elements provided the opportunity to investigate the ages for the LTR elements and families within IBS regions. Many (79%) of these LTR elements are very young (have LTR similarity > 99%) (Table S6). The non-shared LTR elements in these IBS regions are from 12 LTR families that include several very large families as well as 4 families with <10 members (Figure S6B). The small families include only elements with highly similar LTRs while the larger families often include older members (diverged LTRs) as well as a subset with highly similar LTRs, which may suggest the potential for on-going movement in these families.

**Figure 4:**
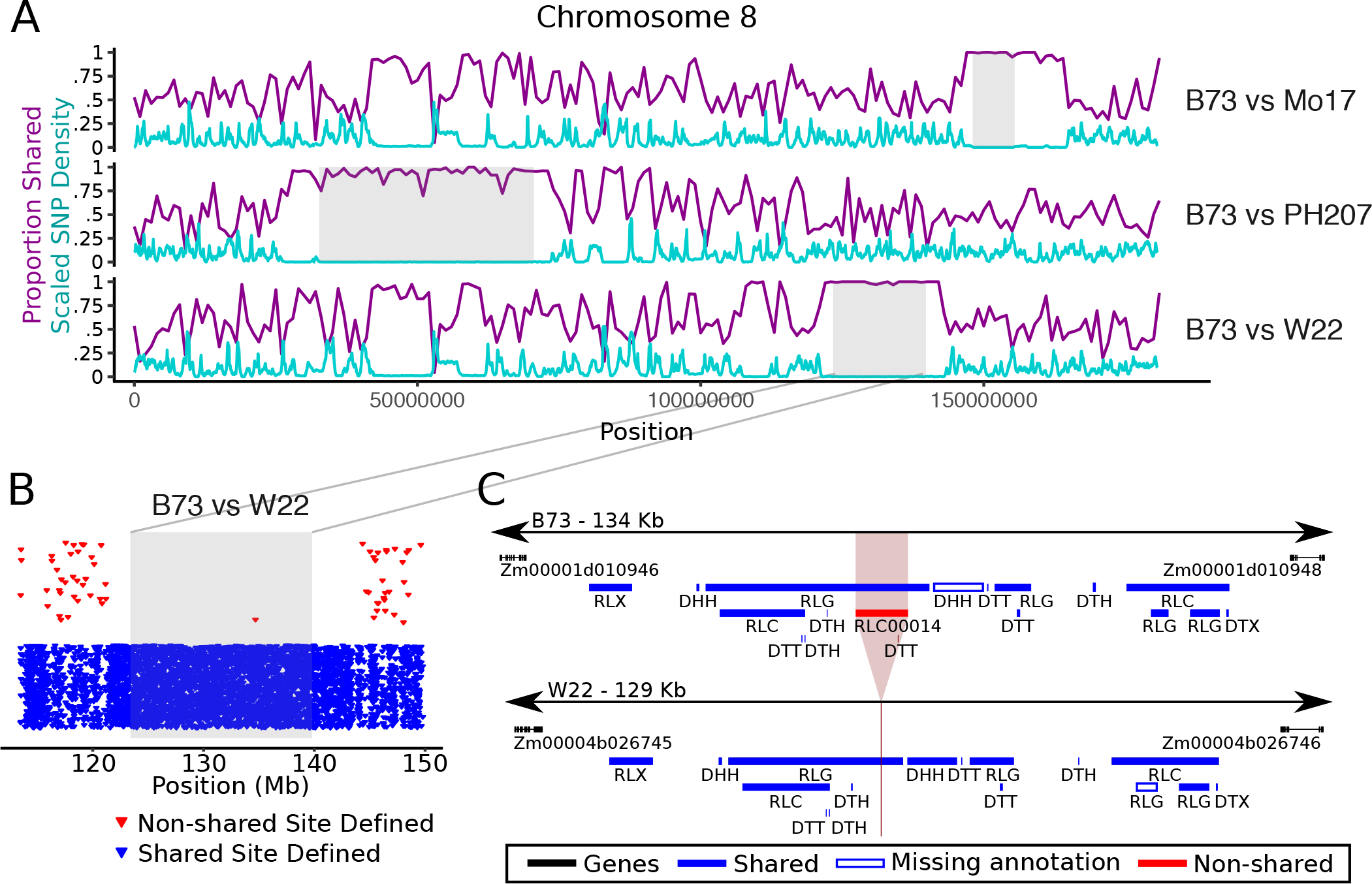
Variable TEs in IBS regions. A. For each contrast against B73, the scaled SNP density and the proportion of variable TEs per 1 Mb bin are plotted. Regions of IBS are shown in gray. B. B73 TEs that are shared and non-shared site defined relative to W22 are plotted for the IBS region and the surrounding 10 Mb. C. There is one new insertion in B73 relative to W22 in the IBS region on chromosome 8, and the region surrounding this insertion is shown.

### Variation in TE content contributes to the extensive gene content variation within maize

While TEs have the potential to influence plant traits through the expression of TE encoded proteins (40, 41), it is largely expected that TEs will influence traits through their effects on genes. Several approaches were used to assess the potential impact of this high level of TE variation on maize genes. We first looked at the proximity of TEs to genes. Over 78% of maize genes have a TE located within the gene or nearby (Figure 5, Table S7). Many of these genes are located near a TE that shows variable presence among the four genomes. Over 7% of maize genes have a variable TE within the annotated transcribed regions, including 1,118 genes containing variable LTRs, 2,384 genes containing variable TIRs, and 176 genes containing variable Helitrons. Surprisingly, over half of all genes have a variable TE within 5 kb upstream of the gene, with 31% near variable LTRs, 30% near variable TIRs, and 10% near variable Helitrons.

**Figure 5:**
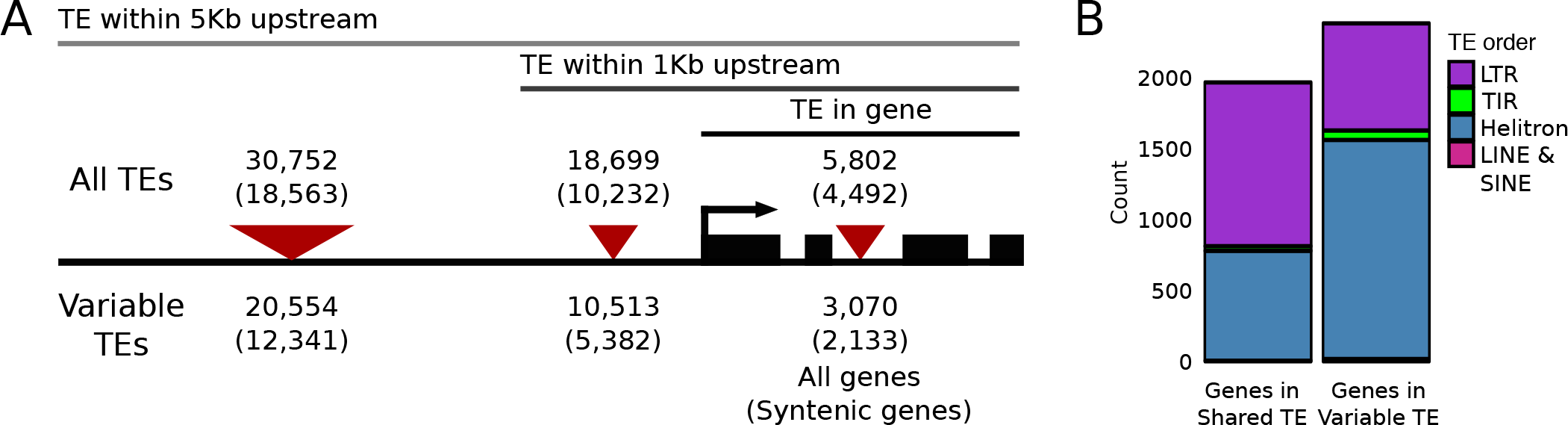
TE contributions to variable gene content in the B73 genome. A. The number of all genes (and syntenic genes) that are overlapping or downstream of any TE (top) or a variable B73 TE (bottom), with the number of syntenic genes listed in parenthesis. TE orders contributing to this distribution are listed in Table S6. B. The 4,344 cases where a non-syntenic gene is completely within a TE are classified based on the TE order and TE variability status.

As we examined the relationship between polymorphic TEs and genes we noted that a substantial number of the polymorphic TEs actually include an annotated gene within the TE. Maize genes (B73v4) were classified as syntenic or non-syntenic based on whether they contained co-linear orthologs in sorghum and other species (37). One of the filters used for the TE annotations included rejecting putative TEs that included a syntenic gene so this analysis was focused on the non-syntenic genes. There are 4,344 non-syntenic B73 genes located within TEs (Figure 5B, Table S8) and 54.8% of these TEs are variable among genotypes. The vast majority of these variable genes were contributed by Helitrons (1,543 genes) and LTRs (754 genes). Presence-absence variation for B73 genes has been noted in all other assembled genomes (29, 32, 34, 35, 42, 43). A comparison of the B73 and W22 genomes identified 6,440 non-syntenic genes that are present in B73 but missing in W22 (34). We find that 20% of these are within non-shared TEs. This highlights the potential for TEs to drive variation in gene content among maize genomes.

The non-syntenic genes located within these variable TEs were further assessed for putative functions and expression. Only a subset (1,277 of 2,380) of the genes have Gramene functional annotations and as expected, some of these were annotated as retrotransposon-like (36 genes), polymerases (27 genes) or helicases (22 genes). However, there were also examples of many annotations that were not related to putative transposon functions such as phytochromes or sucrose synthases. In many cases, the same functional annotation is assigned to multiple genes contributed by the same Helitron family, consistent with one gene capture event preceding TE amplification (Table S8). Additionally, non-syntenic genes located in variable TEs were expressed at low levels and in few tissues. The non-syntenic genes that are located within shared TEs or not within TEs show similarly low levels of expression, which contrasts with syntenic genes which are often expressed in all tissues surveyed. The majority (60%) of the genes in variable TEs are never expressed, show clear reductions in maximum expression values across libraries, and show high levels of CG and CHG methylation (Figure S7). However, the non-syntenic genes not located within TEs tend to have lower levels of CG and CHG methylation. Although the majority of variable genes carried by TEs likely represent inactive pseudogenes or gene fragments, a subset are expressed and have the potential to contribute to phenotypic differences among these lines.

## Discussion

Maize has been a model system for the study of TEs since they were first discovered in the species by Barbra McClintock (1, 44). Active TEs have been used as insertional mutagens to generate functional genomics resources (2, 3, 45), but there are also many examples in which naturally occurring TE polymorphisms have been shown to alter phenotypes. For example, in maize, molecular analyses of QTL involved in domestication or movement from tropical to temperate environments have revealed that TE polymorphisms are responsible for regulatory changes at key loci (26, 46–49).

The variation among individuals within a species is a key aspect of genetics. In the genomics era, researchers typically start from a reference genome and later consider the variable components, often utilizing short-read based resequencing data. This has led to many fruitful studies of genetic variation. However, this approach can be difficult to implement for transposable elements and other repetitive sequences. Prior studies of several loci had suggested high levels of variation in TE content among maize haplotypes (21, 22, 50), but this has not been characterized at the genomic level using whole genome assemblies. In this study we used a novel approach to assess the shared and non-shared nature of TEs within collinear homologous blocks of four assembled maize genomes. The four maize genomes compared in this study represent temperate-adapted maize inbreds (6, 29, 34, 35) and only represent a small fraction of maize diversity (51). These four assemblies are each ~2 Gb in size and we find ~0.45 Gb of TEs that are shared among all four lines. There is another 1.6 Gb of variable TEs that are present in some lines and missing in others. This highlights the massive variability in genome structure of these individuals in a relatively narrow germplasm pool within the same species that is revealed by comparisons of whole-genome assemblies.

Prior studies of TE variation across rice (24) and Arabidopsis (23) accessions used short-read data to identify non-reference TEs with defined insertion sites, which are comparable to our non-shared site defined calls. The analysis of 3,000 rice genomes identified 53,262 non-reference TE insertions (24) and in Arabidopsis 2,835 insertions were identified in a panel of 211 accessions (13). In this study of just four maize genotypes we found 12,620 site-defined TEs present in at least one of three genotypes (W22, Mo17, and PH207), but absent in B73, and a total of 208,535 non-B73 TEs (site defined and non site defined) identified in our full method. Importantly, discovery of these TE polymorphisms lacking homologous flanking sequences requires genome assemblies or long reads as short-read data is not sufficient for the discovery and proper placement of these polymorphisms.

The finding that the majority of non-shared TEs could not be resolved to a specific insertion site due to a lack of homology for flanking sequence was surprising and points to the variety of mechanisms that could result in variable TEs. In some cases these represent TE insertions into a sequence that is not shared between the genomes. In other cases, these represent ancestral TE insertions that have been removed by a larger deletion from one haplotype. This could be due to imprecise TE excision events for some TIR elements or due to larger deletions, which are common and allow for genomes to eliminate intergenic TE sequences (52, 53). This observation that many non-shared TEs also lack homology for flanking sequences creates challenges for studying the consequences of TE polymorphisms on surrounding chromatin, although the subset of site defined TE polymorphisms can be used to study many of the impacts of TE insertions.

The vast majority of previously detected transposition events in maize are due to activity for a small set of TIR families in very specific germplasm (4, 54). On-going movement of LTR transposons has not been detected in maize, although several recent transposition events were detected in large genetic screens (55). The frequency of transposition events in modern maize inbreds appears to be quite low as very few novel mutations due to transposon insertions have been identified. It is likely that the majority of TE variation noted in this study is due to transposition events that have occurred over thousands to millions of years (56, 57). However, the identification of variable TEs in several large IBS blocks does suggest potential for low levels of transpositional activity in the last few hundred generations for some TIR and LTR families in modern maize inbreds. Importantly, these novel insertions will create novel variation that is not tagged by SNPs and could represent functional variation that is not well captured in GWAS approaches.

Variable TEs have likely driven differences in annotated gene content among maize lines. One of the first studies of haplotype diversity in maize revealed the presence of several non-shared genes, as well as numerous non-shared TEs (22). These non-shared genes were later revealed to be gene fragments contained within a Helitron element (58). Several studies have suggested that Helitrons could be a significant source of transposed gene fragments (35, 59, 60). More broadly, a number of studies have found evidence for widespread presence-absence variation (PAV) of gene sequences among maize inbreds (29, 31, 32, 34, 35, 37, 42). We noted >4,000 non-syntenic maize genes located within annotated TEs, including many Helitrons and LTRs. Over half of these represent TE insertions that are variable among the four haplotypes, which will lead to PAV for the genic sequence contained within the TE. The dynamic landscape of TEs among maize genomes leads to high levels of variation for content of gene-like sequences that could have function themselves or could affect the level of expression of related genomic sequences through sRNA mediated influences (61).

The careful classification of shared and variable TEs in maize will create new opportunities to assess the functional consequences that TEs have on genes and genomes. This will enable population level analyses of TE polymorphisms, evaluation of the interplay between TEs and chromatin modifications, and detailed studies of the impact of TE insertions on the expression of nearby genes.

## Methods

### Genome Versions

Results in this manuscript compares whole genome assemblies for four maize genomes: B73 (6), W22 (34), Mo17 (35), and PH207 (29). Analysis was restricted to the assemblies of chromosomes 1 – 10, omitting sequences and annotations assigned to scaffolds.

### TE annotation by nesTEd

NesTEd utilizes several separate programs to annotate different orders of TEs. Briefly, LTR retrotransposons were annotated with a combination of LTRharvest (62) and LTRdigest (63), with iterative removal of identified elements in order to access the often-nested architecture of LTR retrotransposons in maize genomes. TIR and LINE elements were annotated by further refining boundaries of TARGeT searches (64) that used the MTEC TE database (5, 65) as queries. This refinement searched for TIRs and TSDs of an appropriate length for the TE superfamily within 200 bp of either end of the subject match in the reference genome of interest. Helitron elements were annotated using HelitronScanner (66), and SINE elements were annotated using SineFinder (67). As each program searches for different criteria, there is the possibility of individual regions of the genome being annotated as arising from different TEs. To resolve this, we allowed fully-nested copies, where both the start and end positions were found within another annotated TE, but further filtered copies where the span of a TE overlaps the start or end coordinate of another TE. This filtration removed first Helitron, next SINE/LINE, next TIR, and finally LTR, roughly in order of the number and length of structural features used to define each TE order.

Annotation files are available at https://mcstitzer.github.io/maize_TEs. The TE annotation files include metadata including LTR similarity scores for full-length LTR retrotransposons as output by LTRharvest, terminal inverted repeat (TIR) lengths and number of mismatches for TIR elements, and target site duplication (TSD) lengths for TIRs, LINEs, SINEs, and soloLTRs. This metadata was used to assess TE age for LTRs (Figure 1).

The TE annotation file was disjoined to resolve nested TE insertions and to create a file where each bp of the genome is assigned to only the element contributing the DNA sequence of that region. This file was used to calculate the length of each TE and the proportion of the genome contributed by each TE order.

TIR elements were filtered to remove annotations which were nested within another TIR element when the disjoined length of the outer element was less than twice the outer element’s TSD length since in many cases this pattern resulted from ambiguity between 2 bp TIR/TSD patterns. The TE annotation was further filtered to remove ~1,800 TEs from each genome where the TE coordinates completely overlapped a gene with synteny to rice or sorghum, suggesting a false positive annotation. Filtered annotation files are available at https://github.com/SNAnderson/maizeTE_variation.

### Identification of collinear and homologous genes

A cross-reference of homologous maize genes between assemblies was produced using multiple complementary approaches used in an iterative fashion. First, a local version of the SynMap pipeline (68), was used in order to identify stretches of collinear genes in pairwise comparisons of genomes. This pipeline used the LAST aligner version 963 (69) to identify hits between CDS sequence from each genome and then incorporates DAGchainer to identify “chains” of collinear hits. The Lastal algorithm was run using default parameters, however hits were later filtered to have a c-score of 0.10 before being supplied to DAGchainer (70). DAGchainer was ran using an e-value cutoff of 0.05, allowing a maximum distance of 10 genes between matches (−D), requiring a minimum of 12 aligned pairs per chain (−A), and a gap distance of 7 (−g).

The Nucmer alignment algorithm within MUMmer version 3.32 (71) was then used to perform whole-genome alignments between homologous chromosomes of each assembly in pairwise fashion (−c 5,000). The ‘show-coords’ command was used to filter any alignments not included in the longest ascending subset (−g flag) and structural gene annotation files are used to identify genic positions in the alignments. The Nucmer-based gene assignments were cross-referenced with the assignments obtained using the SynMap pipeline. Any gene assignment unique to Nucmer was required to be no further than 3 genes upstream and downstream from the nearest SynMap based gene assignment. This allowed genes that are split across multiple gene models, genes affected by local rearrangements, and genes missed by the SynMap approach to be recovered.

A third approach using the OrthoFinder clustering algorithm version 2.2.7 (72) was used to identify homologous genes. OrthoFinder was run in manual mode by first performing blastp (73) searches across genomes requiring an e-value of 1e-3. The clustering algorithm was then run with the ‘-og’ option to output groups. Each orthogroup cluster was scanned to identify genes from different genotypes that were located on homologous chromosomes. Similar to the nucmer-based gene assignments, collinearity between genes meeting these criteria was assessed using the SynMap and Nucmer assignments. Any assignment unique to this method was required to be no further than 8 genes upstream and downstream from the nearest assignment.

### Identification of shared and non-shared TE insertions

Classification of shared and non-shared elements was performed in a pairwise manner for all contrasts by comparing a TE annotation from one genome assembly (query genome) to the sequence of another genome assembly (target genome). The search window for each TE was defined by the closest, non-overlapping genes in the query genome on either side of the TE that has a unique syntelog in the target genome. In cases of multiple syntelogs or ambiguous syntelog assignment (from one of three methods described above), the outermost gene (relative to the TE) with a unique syntelog assignment was used to define the search window.

For step 1 of the comparison method, a left 400 bp flank tag centered at the start coordinate and a right 400 bp flank tag centering at the end coordinate were first extracted for each TE in one genome (i.e., query genome). These flank tags were then mapped to the another genome (i.e., target genome) using BWA-MEM (74) in paired-end mode. Only cases where the flank tags mapped completely within the search window defined by homologous collinear genes were kept for further characterization. TEs were defined as shared site defined (TE.4, blue in Figure 1A) when at least one flank tag aligned uniquely in the search window with > 90% sequence identity over 90% of the length of the flank tag. TEs were called non shared site defined (TE.5, red) when both flank tags mapped uniquely to the window but both were soft-clipped to the region corresponding to the sequence outside of the TE insertion (with the absolute distance between the left and right outer sequence less than twice the TSD length for the superfamily). TEs nested inside of non shared site defined elements were classified as non shared. In many cases where we called non shared site defined insertions, the left and right flank tags overlapped by the expected TSD length. For 73% of cases in the B73 to W22 contrast, identical sequences were found flanking the B73 TE and at the predicted insertion site in W22, providing further support of a B73-specific insertion (Table S9).

Step 2 of our method utilized full alignment of sequences extracted from query and target windows. LASTZ version 1.03.02 (75) was used to align query and target sequences for each window, with homology blocks defined when sequences shared >80% identity with the ‘--gfextend’, ‘--chain’ and ‘--gapped’ options used. Overlapping alignment blocks were merged and the proportion of each TE that could be aligned between the query and target windows was determined. Elements which were not previously called as site defined (shared or non-shared) were defined as shared (TE.1, dark blue) when coverage was > 90% and non shared (TE.2, dark red) when coverage was < 10%. Elements with intermediate coverage remained unresolved (TE.3, gray).

### Identifying missing TE annotations

In many cases, we found that our method identified shared TEs that were annotated in only one genome due to our strict filtering of structurally intact TEs (Figure S1). In order to capture high-confidence missing annotations, BEDTools intersect (76) was used to compare inferred coordinates for shared site defined elements from each contrast to the TE annotation file. Missing annotations were defined in cases where the inferred coordinates did not intersect with any annotated TE but where the absolute difference between the length of the annotated TE in the query genome and the distance between inferred coordinates in the target genome was less than 1% of the annotated TE’s length.

### Non-redundant TE set

A non-redundant TE set was created for all elements that could be resolved as shared or non-shared compared to all other genome assemblies. This set was created in an iterative manner, starting with all B73 TEs that could be resolved compared to W22, Mo17, and PH207. Resolved W22 TEs that were non-shared with B73 and those that were defined as missing annotations in B73 were then added. Mo17 TEs called non-shared with both B73 and W22 and those that were shared with either B73 or W22 but were defined as missing annotations where shared were added. Finally, fully resolved TEs unique to PH207 and those that were shared with any combination of B73, W22, and Mo17 but were defined as missing annotations where present were added. This resulted in a set of 510,533 TEs present in at least one genome (Supplemental dataset X). The source of each TE annotation can be determined by the genome identifier in the TE name, labeled as Zm00001d for B73, Zm00004b for W22, Zm00014a for Mo17, and Zm00008a for PH207. Disjoined TE length and LTR similarity for the non-redundant TE set were extracted from the genome which was the source of the annotation. TEs were defined as shared when present in all four genomes and as variable when called non-shared with at least one other genome. For family and superfamily-level analyses, the non-redundant TE set was used to summarize TE variability across genomes. This means that in many cases the number of members analyzed for a given family is often less than the total number of annotated elements. Families unique to one genome were required to have annotated copies in only one genome and resolved members in only one genome. Resolved and annotated members in these families are listed in Table S3.

### Characterization of IBS regions

SNPs between B73 and the other three genomes were identified by first aligning these genomes using minimap2 (77). BLAT (78) chain/net tools were then used to process alignment results and build synteny chains and nets. Final SNP and Indel calling was done using Bcftools (79). SNP density for each 1 Mb bin was determined by dividing the total number of SNPs in the window by the number of base pairs in syntenic alignments in the window. Regions with SNP density lower than 0.0005 over at least 5 Mb window were defined as IBS regions. For each comparison between B73 and a contrasting genome (W22, Mo17, or PH207), the inferred coordinates for the outermost shared site defined B73 TEs completely within each IBS block were used to mark the boundary of the IBS region in the contrasting genome.

Genetic distances of IBS regions were calculated using B73 coordinates, a 0.2 cM genetic map (80), and interpolated using a monotonic polynomial function in the R package MonoPoly (81).

### Chromosomal distributions

The distribution of genes and TEs were calculated for 2,106 1 Mb bins across the B73 genome. Exons for isoform 1 were used for calculating gene density and the disjoined TE file was used for TE density. Each feature was assigned to the bin containing the start coordinate from the corresponding gff file. Relative density for annotations was calculated by dividing each bin value by the max bin value across all chromosomes. To calculate TE variability per bin, the full B73 TE annotation file was used to assign TEs to bins based on start coordinates, and TEs were defined as variable when called non-shared in contrast to any other genome.

To assess DNA methylation of the maize genome, B73 was grown and after approximately 6 days shoot tissue was harvested and frozen in liquid nitrogen for use in DNA isolation. Genomic DNA (1μg) was sheared to a size of 200-300bp and the KAPA library preparation kit (KK8232) was used to construct a whole-genome bisulfite sequencing library. The resulting library, which has a size between 250bp and 450bp, was treated with bisulfite sodium so that unmethylated cytosine could be converted to uracil using Zymo EZ DNA methylation lightning kit (D5031). The KAPA HiFi HotStart Uracil + (KK2801) was used in the PCR reaction with the following program: 95°C/2min, 8 cycles of 98°C/30s, 60°C/30s, 72°C/4min, and a final extension step at 72°C for 10 min. The library was sequenced using Illumina HiSeq2000 in paired-end mode with 100 cycles. The WGBS data set has been deposited into NCBI under accession numbers XX.

Trim_galore was used to trim adapter sequences and read quality was assessed with the default parameters and paired-end reads mode. Reads that passed quality control were aligned to B73v4 assembly using BSMAP-2.90 (82), allowing up to 5 mismatches and a quality threshold of 20 (−v 5 -q 20). Duplicate reads were detected and removed using picard-tools-1.102 and bamtools. The methylation level for each cytosine using BSMAP tools was determined. The methylation ratio for 1 Mb non-overlapping sliding windows across the B73v4 genome in all three sequence contexts (CG, CHG, and CHH) was calculated (#C/(#C+#T)).

Chromosomal bins were assigned to quartiles based on gene density and CHG methylation levels and assessed for TE variability among those bins. To assess sub-genome bias in TE variability, bedtools intersect was used to find TEs completely within regions previously assigned as maize1 and maize2 (37). TEs outside of these syntenic blocks are shown as NA.

### Relationship between genes and TEs

B73 genes not present in W22 or sorghum were obtained from previously published results (34). All genome-wide comparisons between genes and TEs were performed using B73 gene and TE annotations. Gene annotations were extracted from the full B73 gff file (6), and coordinates include UTR sequences for the longest defined isoform of each gene. Bedtools intersect (version 2.17.0) was used to find TE-gene overlaps, and bedtools closest was used to find the closest TE of each category. The comparison of genes in TEs included only genes where the full annotated sequence was within the coordinates of a TE, and in cases where a gene was annotated within a nested TE, only the inner-most TE that fully contained the gene was used for analysis. The functional annotations for B73 genes was downloaded from Gramene (ftp://ftp.gramene.org/pub/gramene/CURRENT_RELEASE/gff3/zea_mays/gene_function/B73v4.gene_function.txt) on 8-Nov-2018.

### Expression analysis

To assess expression of genes and TE families across tissues, RNA-seq data from a previous study including 23 tissues of B73 plants were analyzed (39). Gene expression values were downloaded from maizegdb.org (walley_fpkm.txt uploaded 20-Jun-2018) on 19-Jan-2019. Genes were considered expressed in every tissue with an fpkm value of at least 1. Per-family TE expression as assessed from the raw reads as in (83). Briefly, reads were trimmed using cutadapt v.1.8.1, mapped to the B73 genome using bowtie2 using the options -g 20 -i 5 -I 60,000. A modified disjoined TE annotation file was created by using bedtools subtract to mask exons from annotated TEs and then gene annotations were added to the gff file. HTseq-count was used to intersect mapped reads with TE and gene annotations and the SAM output was parsed using a custom script where TE family counts include both reads that map uniquely to a single TE or when multi-mapped but hit only a single TE family. Any read mapped to an annotated gene was not counted towards any TE. Since the length of the expressed fragment cannot be easily estimated for TE families, expression was summarized as reads per million where total reads includes TE family reads and reads mapping uniquely to a gene. TE families are considered expressed when at least one library has an RPM value greater than 1. Per-family expression was plotted as log2(1 + Reads Per Million mapped reads).

## Acknowledgements

We are grateful to Peter Crisp for helpful suggestions on this manuscript. S.N.A., C.D.H, and N.M.S. are supported by a grant from USDA-NIFA (2016-67013-24747). Research on this project was supported by grants from the NSF Plant Genome Research Program (IOS-1238014 for J.R.I and M.C.S.; IOS-1546727 for A.B.B. and C.N.H.; IOS-1546899 for P.Z. and N.M.S.) J.M.N. is supported by a Hatch grant from the MInnesota Agricultural Experiment Station (MIN 71-068). A.B.B. is supported by the DuPont Pioneer Bill Kuhn Honorary Fellowship and the University of Minnesota MnDRIVE Global Food Ventures Graduate Fellowship. The Minnesota Supercomputing Institute (MSI) at the University of Minnesota provided computational resources that contributed to this research.

