## Supplemental figures S1-S7 for "Transposable elements contribute to dynamic genome content in maize"

### Figure S1

A

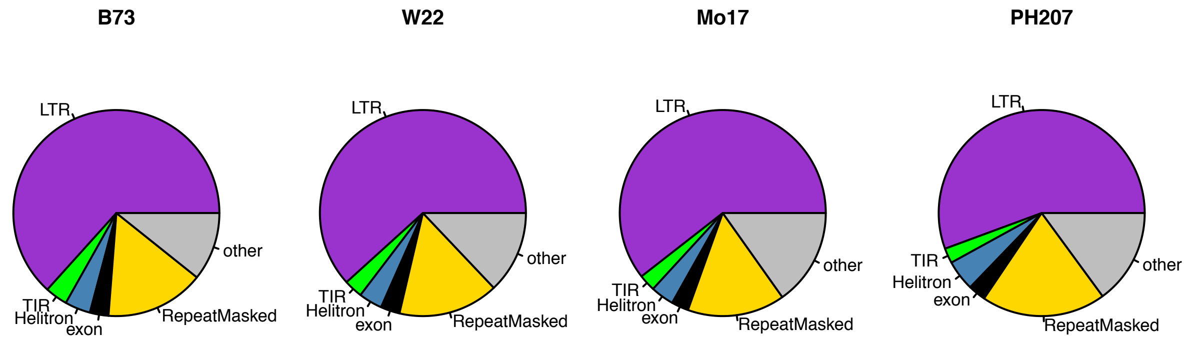


B

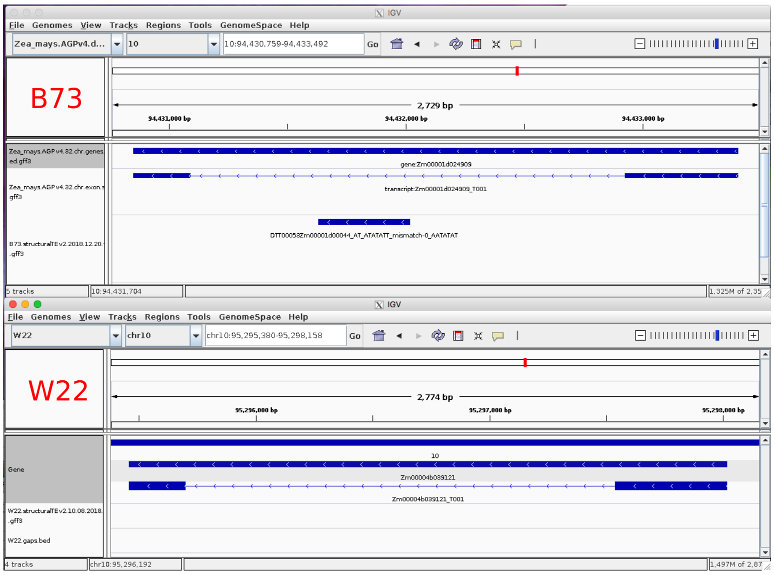


C

```
Zm00004b039121      ACCACTATACATATTGGATTGTTTCTTCACCACA--ATATACAAGAAAATATTAATATCT      1435
Zm00001d024909      ACCACTATATATATTGGATTGTTTCTTCACCACTATATATACAAGAAAATATTAATATCT      1439
DTT00053            --------ATATATTGGATTGTTTCTTCACCACTATATATACAAGAAAATATTAATATCT      52
                            * ***********************   ************************
                            7bp TIR

Zm00004b039121      GCAGTACATATTTGGTGTCATTAAATATGTCTTTTGAAACTATTTTCATAATAAACATAT      1495
Zm00001d024909      GCAGTACATATTTAGTGTCATTAAATATGTCTTTTGAAACTATTTTCATAATAAACATAT      1499
DTT00053            GCAGTACATATTTAGTGTCATTAAATATGTCTTTTGAAACTATTTTCATAATAAACATAT      112
                    ************* **********************************************

Zm00004b039121      TTGAAGATACAGATATTTCAAATATTTTTTACGAATCTAATCAAATATGAGAAATTTTGA      1555
Zm00001d024909      TTGAAGATACATGTATTGCAAATATTTTTTACGAATCTAATCAAATATGAGAAATTTTGA      1559
DTT00053            TTGAAGATACATGTATTGCAAATATTTTTTACGAATCTAATCAAATATGAGAAATTTTGA      172
                    ***********  **** ******************************************

Zm00004b039121      CTGACATGTATGACCATACTATCAATTATTTTAGGACAGAGGCAGGGAGTGCGCATTTGT      1615
Zm00001d024909      CTGACATGTATGACCATACTATCAATTATTTTAGGACAGAGGCATGGAGTGCGCATTTGT      1619
DTT00053            CTGACATGTATGACCATACTATCAATTATTTTAGGACAGAGGCATGGAGTGCGCATTTGT      232
                    ******************************************** ***************

Zm00004b039121      ATGGTCGAAATCGATCAATTGTAACCATATGCGTGTACG---TTGGTACGCCCACTGATG      1672
Zm00001d024909      ATGGTCGAAATCGATCAATTGTAACCATATATGCATGTACGTTTGGTACGCCCACTGATG      1679
DTT00053            ATGGTCGAAATCGATCAATTGTAACCATATATGCATGTACGTTTGGTACGCCCACTGATG      292
                    ******************************  *  *      ******************

Zm00004b039121      TATCTACCTGGTTAATTAATTAGATATATGACCTAGCTTGTCGTCTGATTGTTATGATTA      1732
Zm00001d024909      TATCTACCTGG----TTAATTAATTAGATGACCTAGCTTGTCGTCTGATTGTTATGATTA      1735
DTT00053            TATCTACCTGG----TTAATTAATTAGATGACCTAGCTTGTCGTCTGATTGTTATGATTA      348
                    ***********    *******  ** *********************************

Zm00004b039121      AAGAACCAAAAAGTCTACTCAGCTCAAAACCCAAATATATATGTGTCAAACAACTCCCAT      1792
Zm00001d024909      AAGAACCAAAAAGTCTACTCAGCTCAAAACCCAAATATATATGTGTCAAACAACTCCCAT      1795
DTT00053            AAGAACCAAAAAGTCTACTCAGCTCAAAACCCAAATATAT--------------------      388
                    ****************************************
                                                     7bp TIR
```

Figure S1: Summary of StructuralTEv2 annotations for four maize genomes. A. The proportion of maize assemblies annotated as LTR (purple), TIR (green), Helitron (blue), or exons (black). The yellow section marks additional repeat sequences masked by Repeat Masker. B. Example locus showing a TIR element annotated within the intron of gene Zm00001d024909 in B73 where no TE is annotated in the intron of the corresponding W22 gene (Zm00004b039121) but the region is highly similar in sequence. The alignment of both genes and the annotated TE reveal a 1 bp SNP in the TIR of the W22 TE, causing the TE to fail our stringent filtering criteria.

### Figure S2

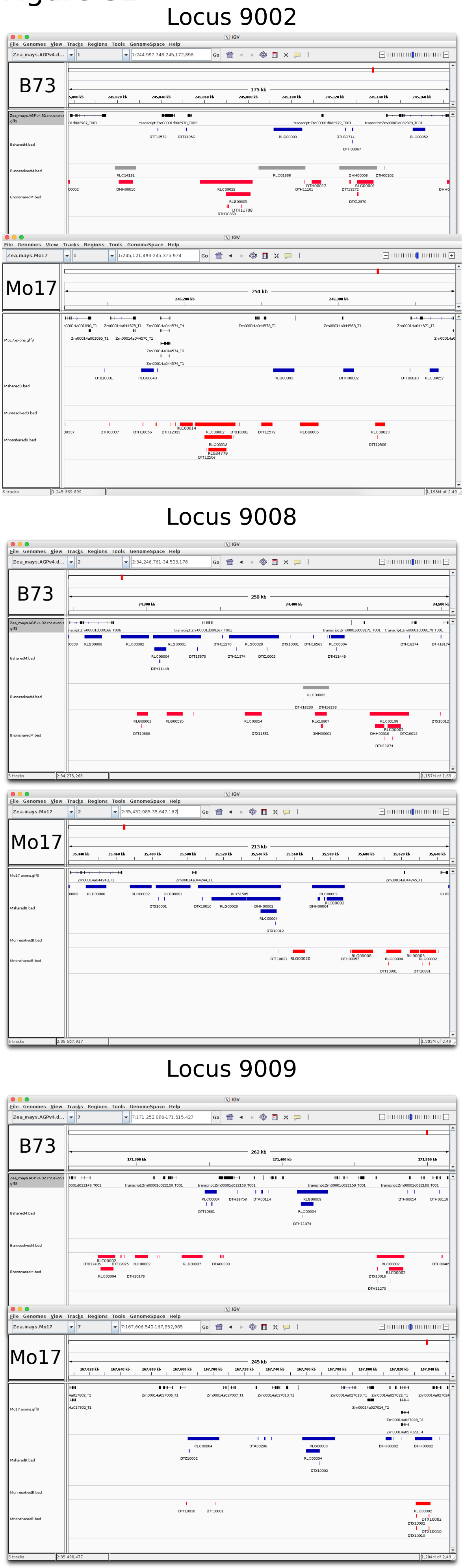


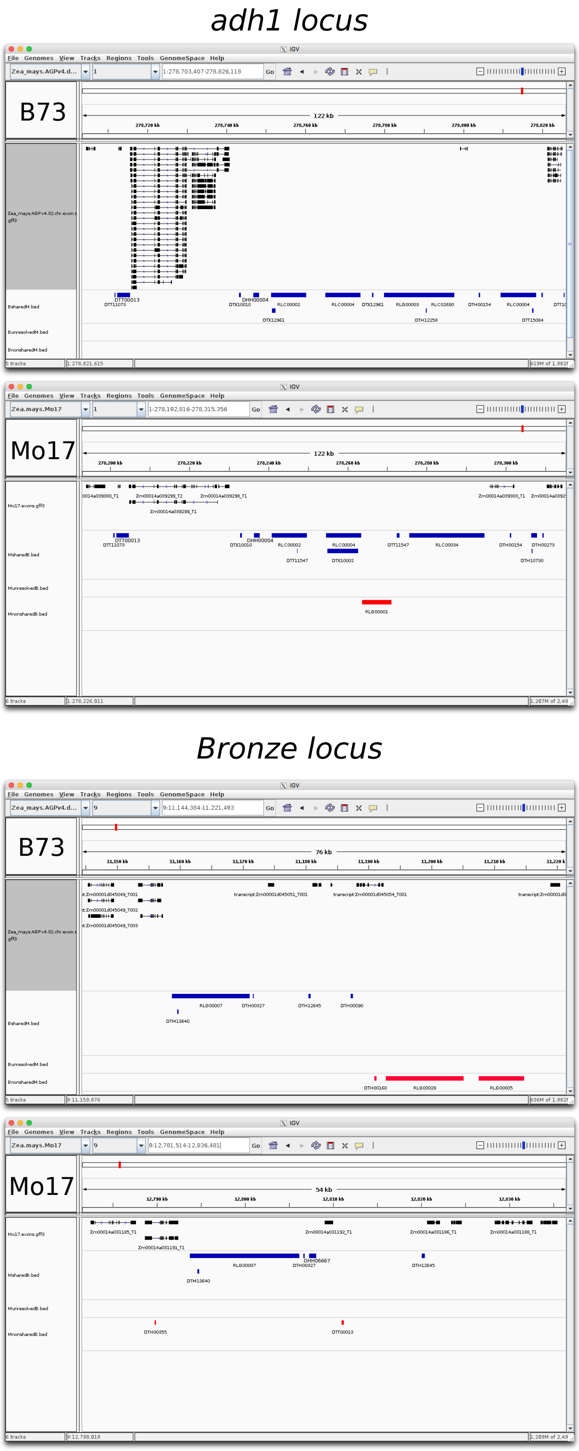


Figure S2: Validation of the classification of shared and non-shared TEs based on comparison to manually annotated genomic regions of B73 and Mo17 (Brunner et al. 2005). Genes (black) and TEs (shared in blue, nonshared in red, unresolved in gray) are visualized. Coordinates shown are limited to the region defined by the outermost shared genes, corresponding to genes A-F for locus 9002, A-E for locus 9008, A-P for locus 9009, A-F for adh1, and A-M for bronze. TEs are labeled by family ID. The genome-wide classifications in this work are in strong agreement with TE polymorphisms noted in the manual annotation of Brunner et al. (2005).

### Figure S3

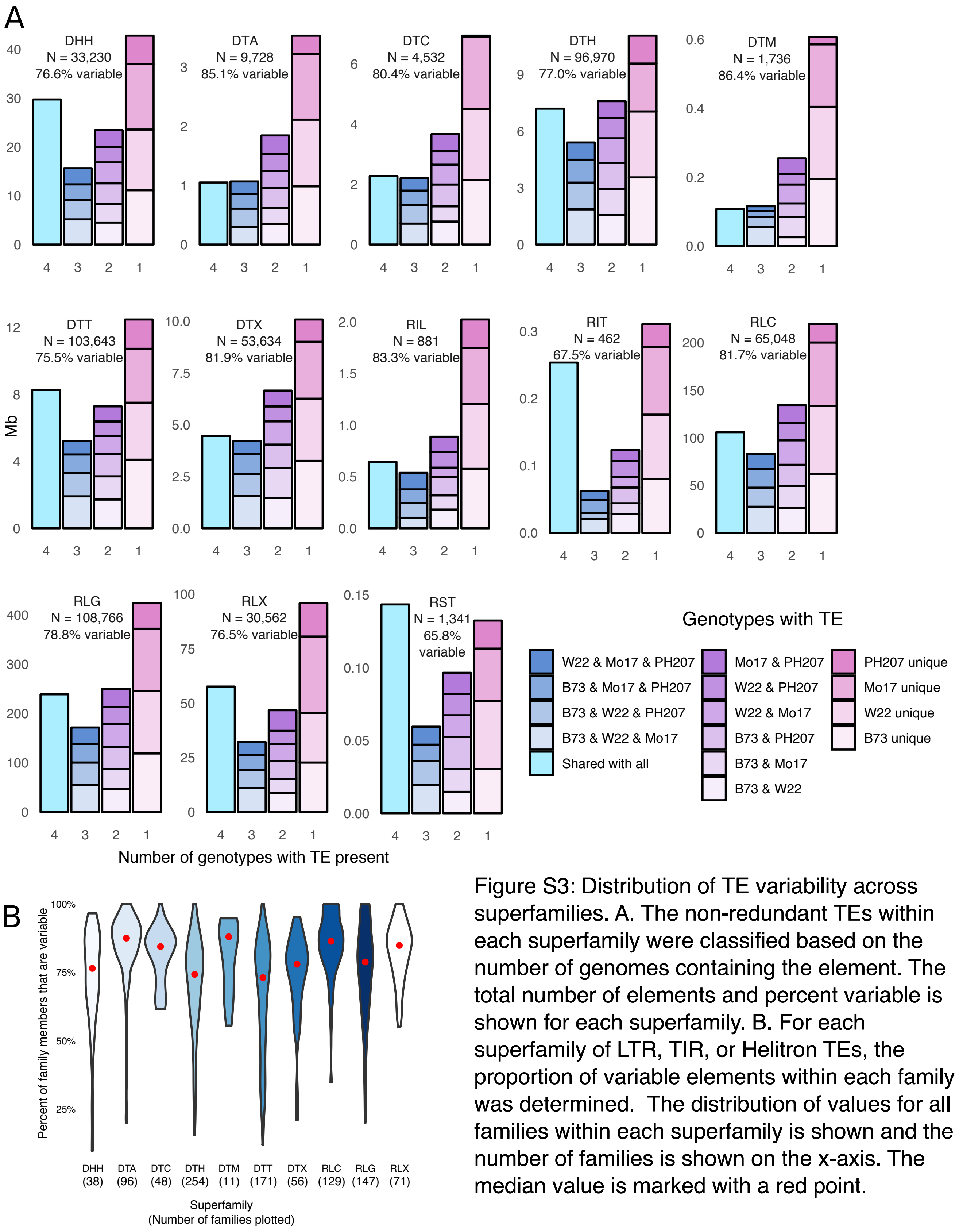


Figure S3: Distribution of TE variability across superfamilies. A. The non-redundant TEs within each superfamily were classified based on the number of genomes containing the element. The total number of elements and percent variable is shown for each superfamily. B. For each superfamily of LTR, TIR, or Helitron TEs, the proportion of variable elements within each family was determined. The distribution of values for all families within each superfamily is shown and the number of families is shown on the x-axis. The median value is marked with a red point.

Figure S4

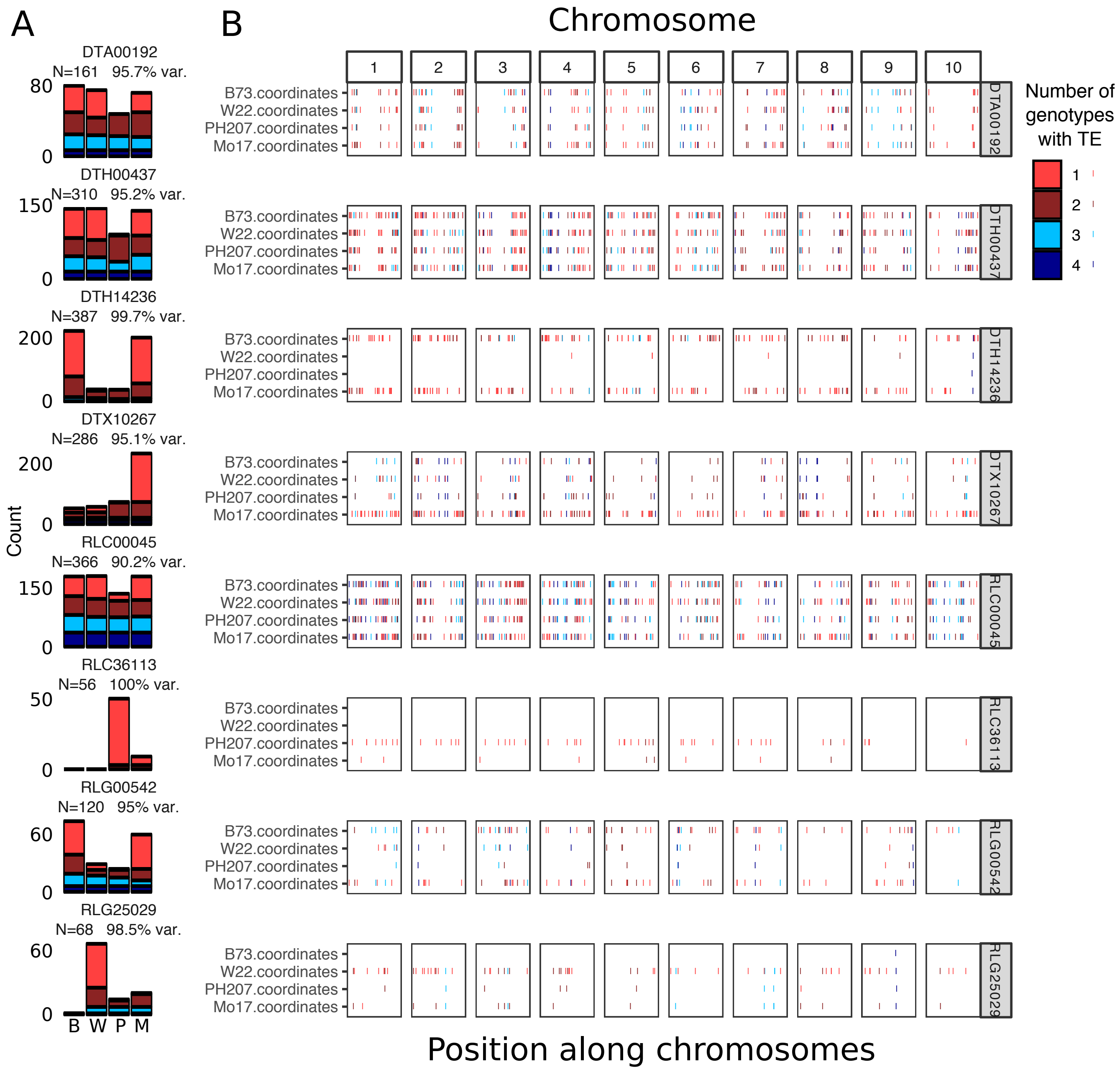


Figure S4: Genome distributions for highly variable TE families. A. For a subset of families with high variability the number of elements in each family and level of sharing is plotted, with the number of members and percent variable listed. B. Chromosomal distributions across genomes for elements in families shown in A. Colors indicate the number of genomes containing each TE.

Figure S5

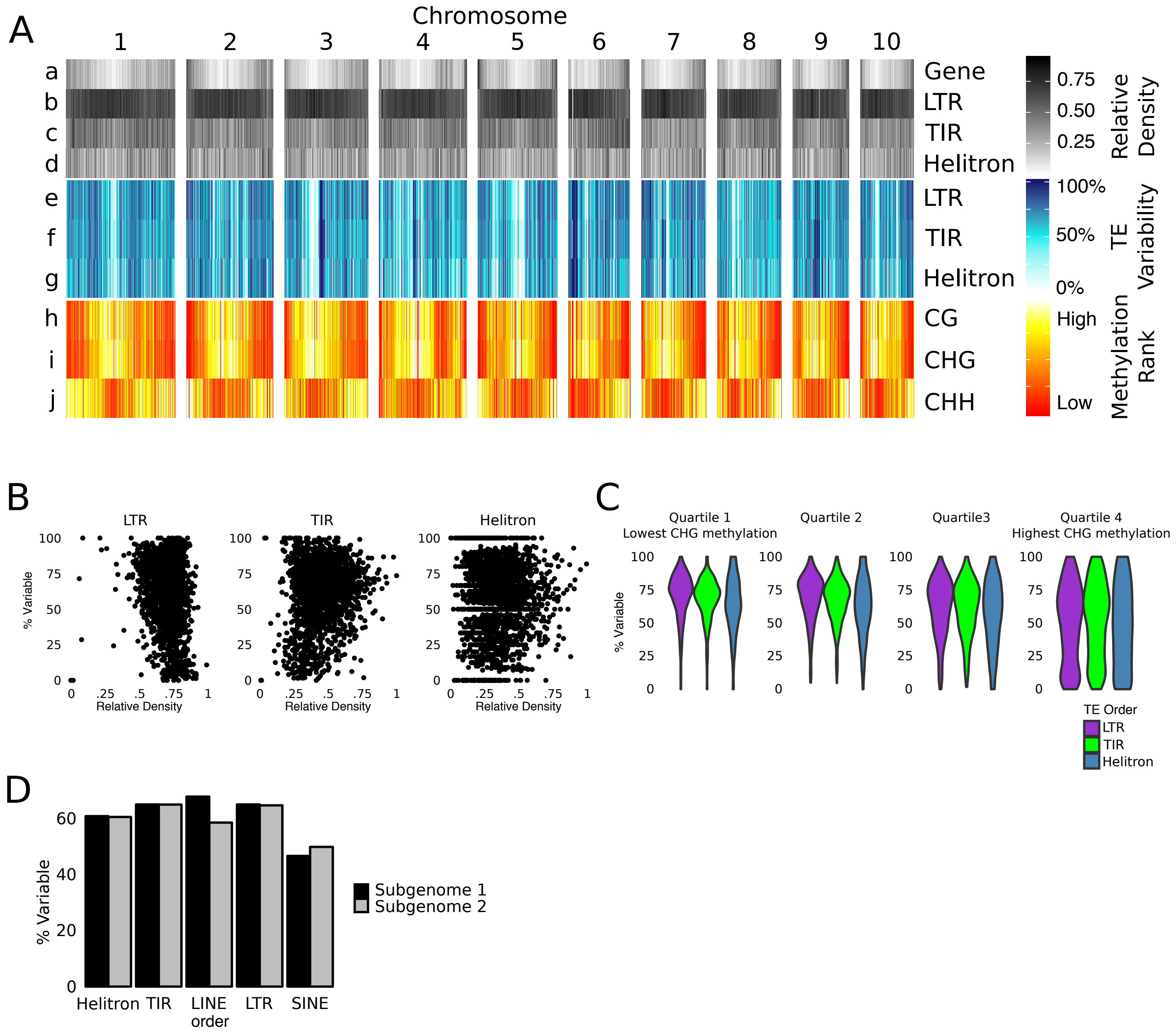


Figure S5: Genome-wide distribution of TE variability. A. The distribution of gene density, TE variability and DNA methylation in 1 Mb bins along all B73 chromosomes. Tracks a-d show the relative gene density and TE density. Tracks e-g show the proportion of B73 TEs that are variable in each bin for LTRs, TIRs and Helitrons, and tracks h-j show the rank order of methylation in the CG, CHG, and CHH context. B. For each 1Mb bin, the relative density and proportion of variable elements were compared for LTRs, TIRs, and Helitrons, showing no correlation for any TE order. C. The CHG methylation level in each 1 Mb bin was used to splits the bins into four quartiles from lowest to highest methylation density. The proportion of variable TEs within each bin were determined for each superfamily. D. The variability for TEs completely within the two maize sub-genomes is highly similar for all orders of TEs.

### Figure S6

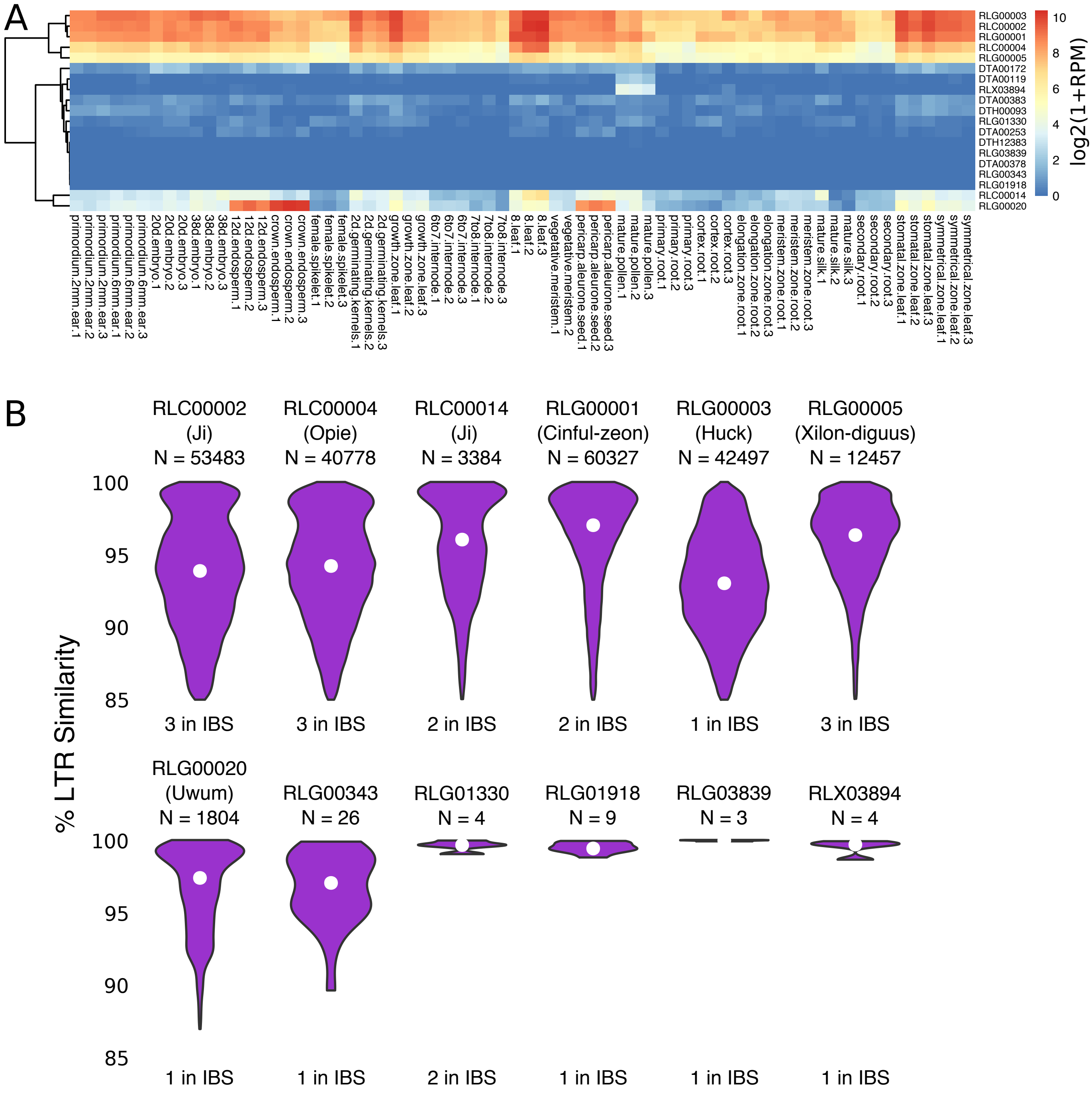


Figure S6: Family attributes for new insertions in IBS regions. A. The expression of each TE family across tissues in B73 (Walley et al. 2016) is plotted, with examples of families expressed across all tissues, some tissues, or never expressed. B. For 12 LTR families with new insertions in IBS reigons, the distribution of LTR similarity scores is plotted. The number of full-length members is listed, and the common name for each family is shown in parenthesis, when applicable. The median LTR similarity for each family is marked with a white point.

Figure S7

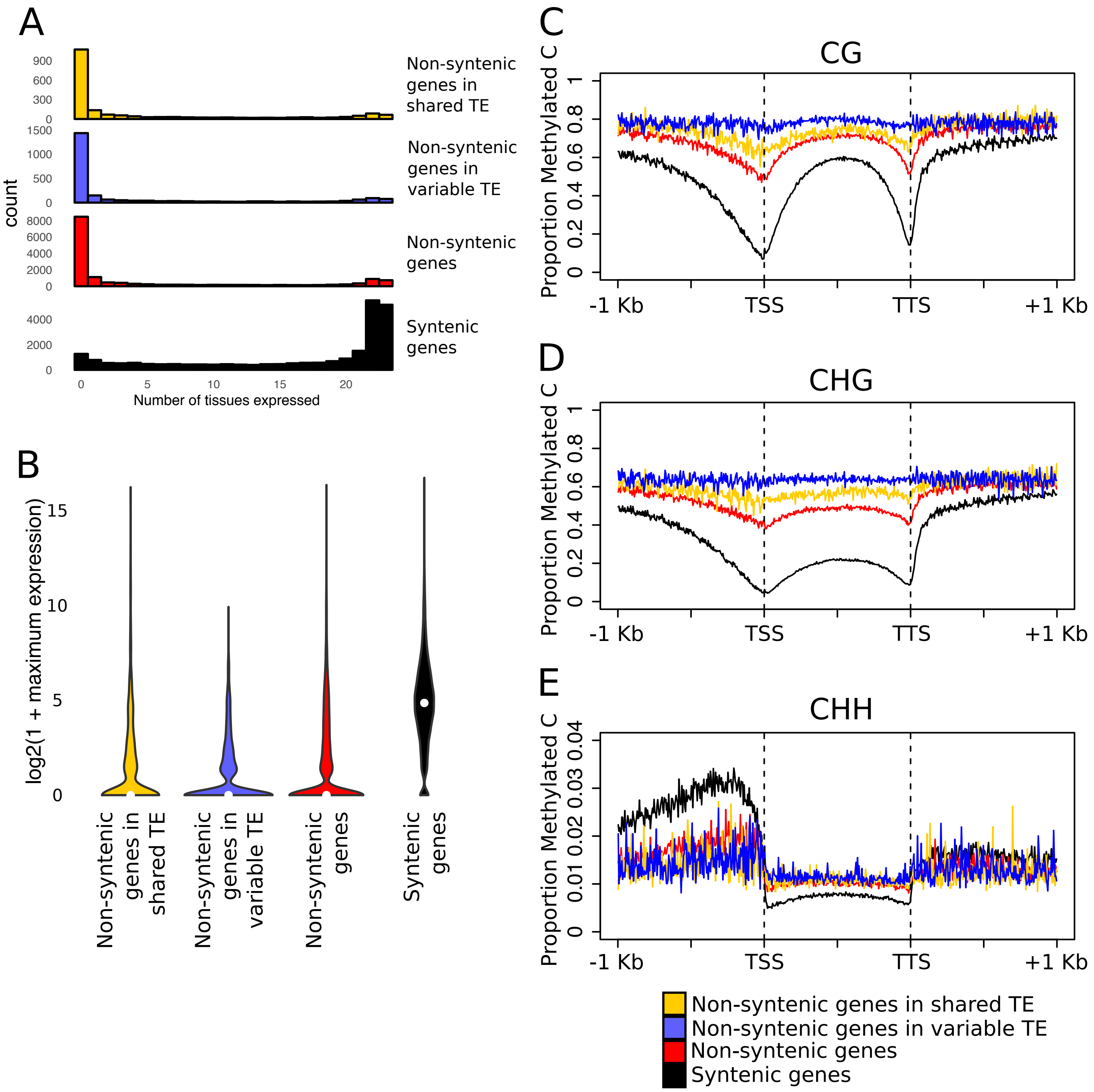


Figure S7: Expression and methylation patterns for genes in TEs. A. The number of tissues with expression of syntenic genes (N=23,922) is compared to non-syntenic genes located in variable TEs (N=2,380), non-syntenic genes in shared TEs (N=1,960), and all other non-syntenic genes (N=15,519) across 23 tissues of B73 (data from Walley et al. 2016). B. The maximum expression value for TEs across B73 tissues is compared for each gene set. White points mark the median value. C-E. DNA methylation within and for the 1 Kb surrounding the genes in each set in the CG (C) CHG (D) and CHH (E) contexts, showing different patterns of methylation for these different gene subsets.
